# RF Channel-Based Adaptive Beamforming for 3D Ultrasound Localization Microscopy

**DOI:** 10.1101/2024.08.02.606290

**Authors:** Georges Chabouh, Baptiste Pialot, Louise Denis, Raphael Dumas, Olivier Couture, Pauline Muleki Seya, François Varray

**Affiliations:** Sorbonne Université, UMR 7371 CNRS, Inserm U1146, Laboratoire d’Imagerie Biomédicale, 15 Rue de l’Ecole de Médecine, 75006 Paris, France; INSA-Lyon, Universite Claude Bernard Lyon 1, CNRS, Inserm, CREATIS UMR 5220, U1294, F-69621, Lyon, France

## Abstract

Ultrasound Localization Microscopy (ULM) has been applied in various preclinical settings and in the clinic to reveal the microvasculature in deep organs. However, most ULM images employ standard Delay-and-Sum (DAS) beamforming. In standard ULM conditions, lengthy acquisition times are required to fully reconstruct small vessels due to the need for spatially isolated microbubbles, resulting in low temporal resolution. When microbubbles are densely packed, localizing a point spread function with significant main and side lobes becomes challenging due to matrix arrays’ low signal-to-noise ratio and spatial resolution. In this work, we applied adaptive beamforming such as high order DAS known as (pDAS), Coherence Factor (CF), Coherence Factor with Gaussian Filtering (CFGF), and statistical interpretation of beamforming (iMAP) to provide a more complete 3D ULM maps *in vitro* and *in vivo* (rat kidney). Specifically, the CF and 1MAP adaptive beamformers achieved higher resolution (32.9 microns and 27.2 microns respectively), as measured by the Fourier Shell Correlation (FSC), compared to the standard DAS beamformer, which had an FSC value of 38.6 microns.

## Introduction

Ultrasound Localization Microscopy (ULM) is a super-resolution ultrasound technique that has gained significant traction for its ability to image deep *in vivo* blood vessels within organs^1^. In humans, it has demonstrated high performance in 2D with both ultrafast acquisitions^2,3^ and the low frame rate line-by-line acquisitions typical of clinical ultrasound scanners^4–6^. Recently, transthoracic ULM has been applied in patients for myocardial microvasculature imaging^7^. With advancements in ultrasound technology, volumetric ULM has become feasible through various techniques such as fully-addressed^8–11^, sparse arrays^12,13^, row-column arrays^14–17^ and multiplexed arrays^18–23^. ULM is based on the localization of single acoustical scatterers^24,25^, generally familiar as clinically approved coated microbubbles^26,27^, and the ability to track them in space and time within the vessels^28,29^. While ULM can reveal capillary-scale vessel sizes^30^, achieving such high resolution comes at a time cost, resulting in shallow temporal resolution^31^. Various approaches have been introduced to overcome this limitation to enhance localization in high microbubble concentrations ^32,33^.

Volumetric ULM offers a promising solution to typical 2D ULM issues such as out-of-plane motion^18^, inaccurate quantifications^34^, and user dependency. However, as discussed by Yan *et al*., matrix probes used for volumetric ultrasound systems have lower spatial resolution and Signal-to-Noise Ratio (SNR) than linear probes due to their small aperture size and element size^23^.

After the acquisition yielding radio-frequency channel data, the ULM processing pipeline consists of various steps, including beamforming, clutter-rejection, sub-wavelength localization of isolated microbubbles, tracking of these super-resolved positions, accumulation, and visualization. Many improvements have been proposed for each of these steps. However, the impact of the beamforming of the radio-frequency data remains sparsely investigated, especially for volumetric (3D) ULM ^23^.

The classical beamformer used before the ULM algorithmic pipeline is usually a simple delay-and-sum (DAS)^35^. Its main benefits are simplicity and processing time, which is especially important in 3D. Several other adaptive beamformers have been introduced in the literature, attempting to shrink the Point Spread Function (PSF) and reduce side lobes. A primary example of an adaptive beamformer is the p-th root compression (pDAS), which involves raising each channel signal to the power of *p*, summing them, and then taking the p-th root of the result. Another way to change the weight for each channel signal before summing them is with minimum variance^36,37^. Additionally, the coherence factor (CF) beamformer reduces the intensity of voxels dominated by noise or incoherent artifacts^38^. Null subtraction imaging has been used to improve the visibility of the microcirculation ^39^.

To date, the only study on the effect of adaptive beamformers for volumetric ULM has been conducted by Yan *et al*.^23^. They introduced a novel adaptive beamformer combining the coherence factor with the variance of the channel signal, termed Coherence Energy to Variance (CV). Their study compared and evaluated DAS beamformer, pDAS, CF, and CV both *in silico* and *in vivo* using rabbit kidneys. The authors demonstrated that the proposed CV beamformer performed significantly better, producing the narrowest main lobes and weakest side lobes.

In this study, we compare various beamformers to investigate their effect on both *in vitro* and *in vivo* volumetric ULM on rat kidneys. We hypothesize that shrinking the PSF and reducing side lobes would benefit ULM as it relies on the precise sub-wavelength localization microbubbles PSF. We evaluated DAS, pDAS, CF, CF with Gaussian Filtering (CFGF), and a statistical interpolation beamformer that maximizes the Maximum A Posteriori (iMAP) distribution of the aperture signals^40^. These adaptive beamformers were exploited by performing clutter-filtering before beamforming, which differs from conventional approaches. We report in this work a better resolution with the adaptive beamformers compared to the standard DAS beamformers for the entire kidney in agreement with previous findings^23^. The data and the proposed adaptive beamforming algorithms optimized for rapid 3D imaging are available as open source for the ultrasound community^41^.

## Materials and Methods

### Characterization of the Point Spread Function (PSF) via in-vitro experiments

he *in vitro* experiment was performed in a 3D-printed box (10 cm ×10 cm ×30 cm) where an acoustic absorber is placed at the bottom. Deionized water at T=25°C was gently poured inside the box to prevent air bubble formation. The water was placed in a glass bottle for a week for total air bubbles dissolution. SonoVue microbubbles (Bracco, Italy) were prepared as mentioned by the manufacturer. They were injected inside the box with a dilution of 1:10000 ratio (in volume) to ensure microbubble separability, a crucial step in super-resolution ultrasound. Finally, a magnetic stirrer is placed at the bottom of the box to prevent microbubble flotation.

### In vivo rat model

Our previous work conducted animal experiments using 3D sensing ULM (sULM) to automatically segment the functional part of the kidney known as the glomeruli^18^. All animal experiments were performed following the ARRIVE guidelines and were approved by the local ethics committee (ethics committee on animal experimentation n°034). The Ministry of Research registered the protocol under number #33913-2021082311153607. Adhering to the 3R principles, the number of animals in our study was minimized. Experiments were performed on eight Wistar male rats (Janvier Labs, Charles River, Envigo) aged between 8 and 12 weeks. In this work, the data from a single rat was used.

### Ultrasound acquisitions

As detailed in Chabouh *et al*.^18^, the left kidney was externalized through an incision in the abdomen (See Figure 1 in^18^ and all the details). Around 5 to 10 mm of acoustic gel is placed between the organ and the probe to eliminate the near field of the transducer. A 25G catheter was placed in the tail vein to perform SonoVue microbubbles injections. The animal was under isoflurane anesthesia (4% for induction, 2.5% for maintenance) with appropriate analgesia (subcutaneous injection of 0.1 mg/kg of buprenorphine 30 minutes before experiments).

**Figure 1.**
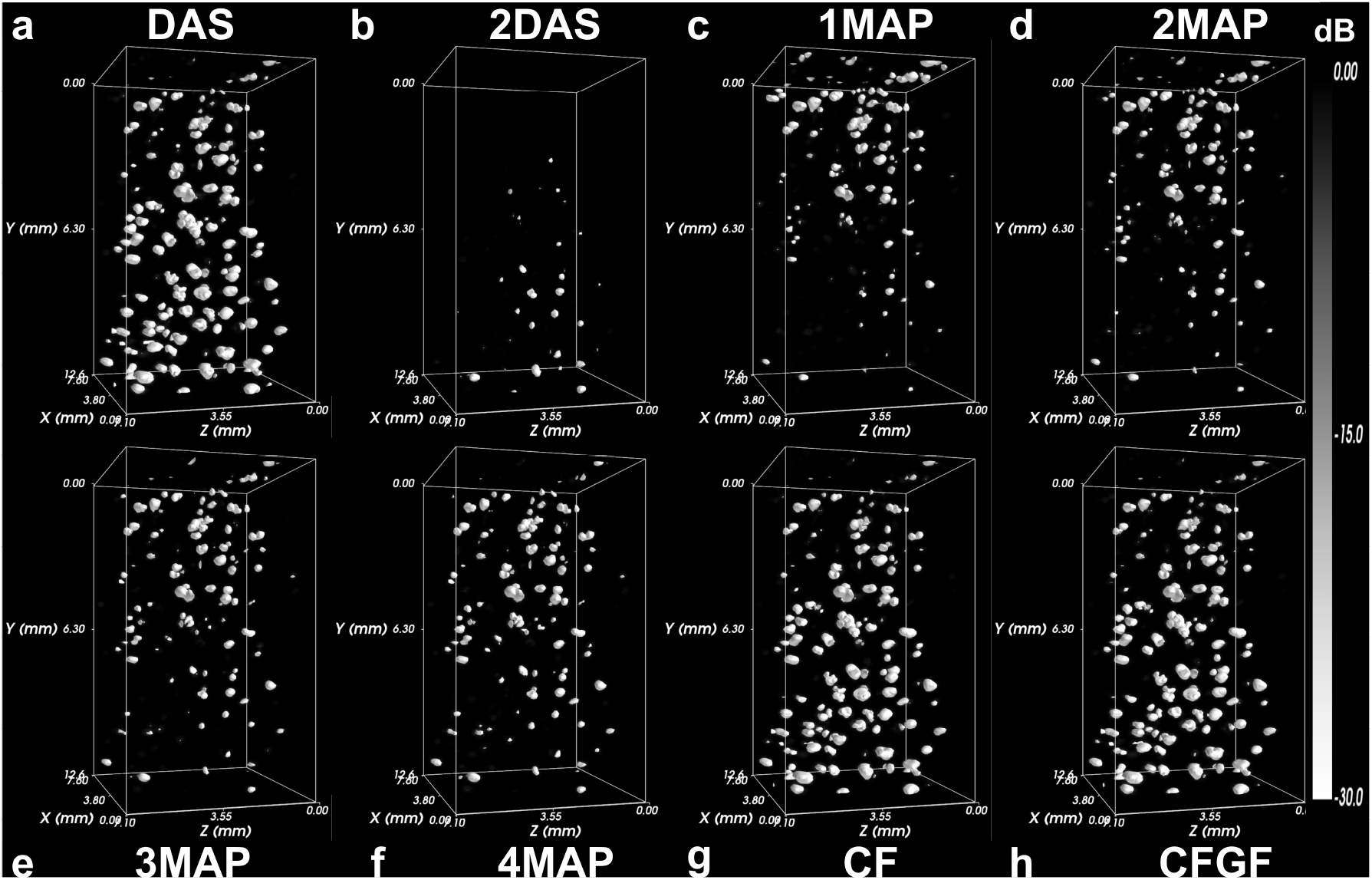
*In-vitro* microbubbles in water renderings with the proposed adaptive beamformers: **a** DAS, **b** 2DAS, **c** 1MAP, **d** 2MAP, **e** 3MAP, **f** 4MAP, **g** CF and **h** CFGF.

Details of the ultrasound acquisition parameters can be found in^18^. It was performed using a 256-channel research ultrasound scanner (Verasonics, Kirkland, USA) and an 8 MHz multiplexed matrix probe (Vermon, France). The probe operates at a central frequency of 7.8 MHz, emitting 2-cycle pulses at a 67% duty cycle. These pulses are driven at 10 volts, resulting in a mechanical index (MI) of 0.1. Five hundred blocks were acquired, with 200 volumes per block and a volume rate of 130 Hz. Each image results from a compounding of 5 plane waves oriented according to *±*5° in the elevation and lateral axis. The multiplexed probe is divided into four panels of 256 elements each. The sequence lasted for 8 minutes, with repeated bolus injections of 50 *μ*L/min every 1 minute. The data were then reconstructed with all the proposed beamformers (see the next section) on a (150, 150, 98.5) *μ*m (*x, y, z*) grid, before reaching the final ULM resolution of (9.85, 9.85, 9.85) *μ*m. To reduce the quantity of acquired data, a 100% bandwidth mode was used on the Vantage system, leading to only two acquisition points per period. Once the IQ signals are generated from the RF data, only a single IQ value is computed per period of raw signals. In other words, the final axial pixel size corresponds to one wavelength.

### Presentation of beamformers

This subsection presents the selected beamforming techniques and recalls their basic principles. Graphical representation can be found in the supplementary Fig. 8. Adaptive beamformers were selected for their proven ability to increase resolution and/or increase contrast, enabling microbubbles to be highlighted in a noisy environment. The beamformers’ speckle-preserving capability was not considered, as it should play a limited role in microbubble localization and tracking.

The variable *s*_*k*_ will denote the delayed IQ signal on the k-th channel for the voxel of index *n*. The DAS output for this voxel is then expressed as:

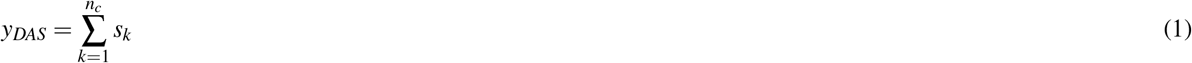

where *n*_*c*_ is the number of channels. The DAS beamforming is conducted based on the *Ultraspy* python package, freely available online^42^. In addition, the various proposed adaptive beamformers presented in this paper can be found in a dedicated git repertory to allow results reproducibility^41^. The proposed implementation can be executed on CPU or GPU. Based on SVD, a clutter filter is also provided (described in section).

#### pDAS

The pDAS adaptive beamformer performs a non-linear p-root compression of delayed IQ signal to reduce the intensity of incoherent voxels between channels. However, the *p* value must be chosen carefully, as nonlinear compression can exclude the areas of the images with the lowest coherence between channels^43^. To be compatible with IQ demodulation, pDAS must be implemented in its baseband version^44^. The first step is to sum the signed p-rooth of the delayed IQs along the channels:

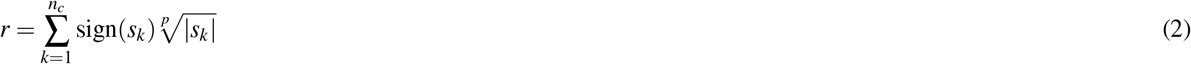

where the sign of the complex signal *s*_*k*_ is defined as:

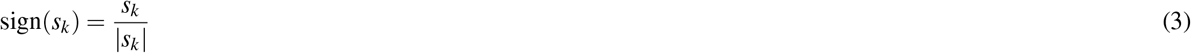

The pDAS output with a restored physical dimension for a given voxel is then:

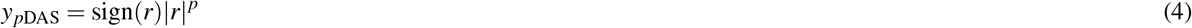

#### CF and CFGF

The CF adaptive beamformer is based on weighting the DAS output by a voxel-wise coherence factor:

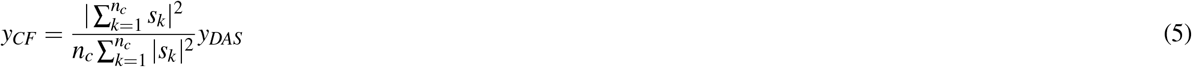

The coherence factor in equation (5) is the ratio between the coherent sum of *s*_*k*_ across channels and its incoherent sum. For a given voxel, *CF* = 1 if its channel signal is purely coherent, and *CF* = 0 if it is purely incoherent. The coherence factor, therefore, reduces the intensity of voxels dominated by noise or incoherent artifacts.

The coherence factor can exhibit significant discontinuities between adjacent pixels, impacting its effectiveness. One way of dealing with this effect is to calculate the coherence factor for all voxels and smooth it across the voxel dimension using a 3D Gaussian kernel^45^. This variation will be called CFGF (Coherence Factor with Gaussian Filtering) and evaluated as a separate beamformer from CF, as it might produce significant differences in the final ULM map, related to the spatial smoothing of the CF factor.

#### iMAP

The iMAP adaptive beamformer models the channel signal for a given voxel as:

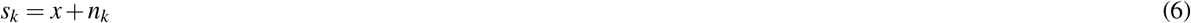

where *x* is the reflectivity of the targets within the voxel and *n*_*k*_ represents interferences such as electronic noise. The quantities *x* and *n*_*k*_ are assumed to be two decorrelated zero-mean Gaussian random variables with standard deviation 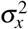 and 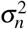, respectively^40^.

From the model in equation (6), it can be shown that the summation of *s*_*k*_ along channels occurring in DAS is equivalent to a maximum likelihood estimate of *x*. This estimate can be improved using a more advanced estimator such as maximum a posteriori (MAP). However, the MAP estimator requires knowledge of 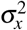 and 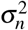, both unknown quantities in practice. To solve this problem, the two standard deviations are iteratively estimated by maximum likelihood (Algorithm 1 in^40^). Once the final iteration is reached and 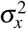 and 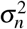 are estimated, the output of the iMAP beamformer is directly obtained.

##### Algorithm 1 iMAP algorithm

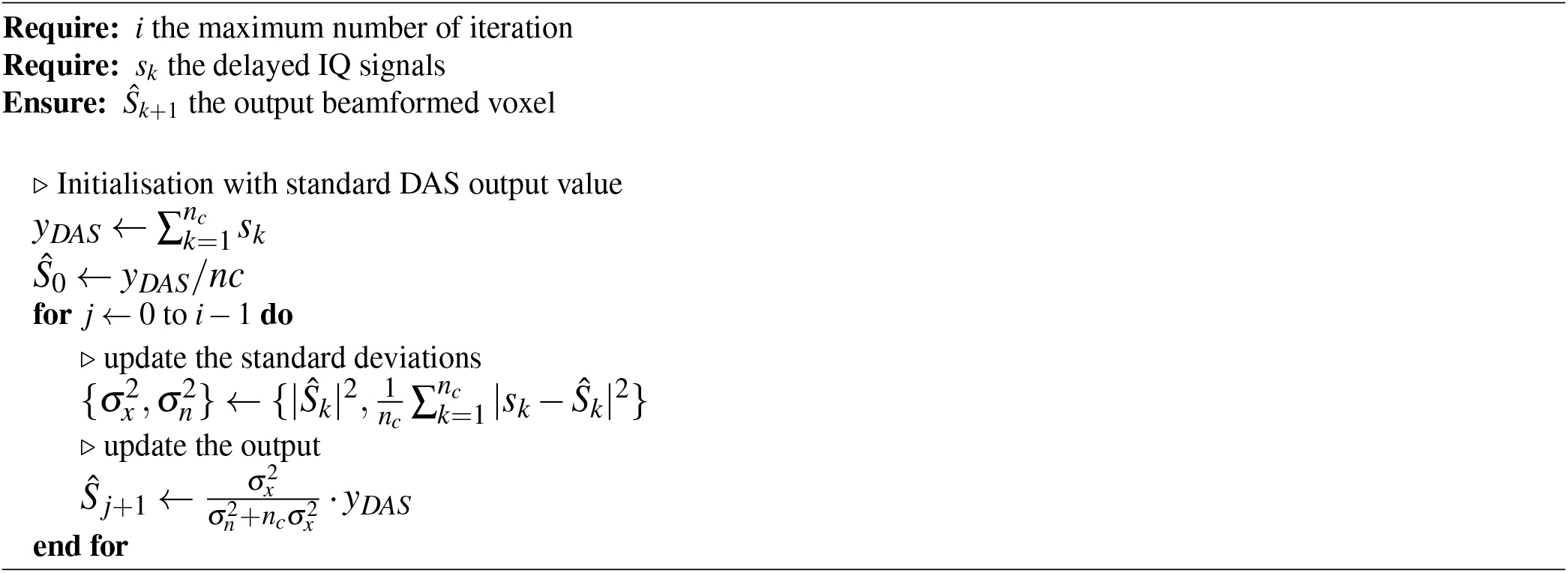

### Clutter filtering and implementation of beamformers

In a standard ULM pipeline, the tissue signal and the motion due to the respiration and heartbeat should be filtered out to enhance the microbubble signal for detection^46^ and tracking^47^ based on microbubble non-linearity^48^. In general, clutter filter techniques are based on spatiotemporal filters, such as singular-value decomposition (SVD), which rank the image components by energy, with higher energy indicating greater coherence in space and time^1,49^. A classical way to apply these filters is after the beamforming step, i.e., after image formation^1^. However, when using adaptive beamforming, it would be of greater importance to evaluate their effect on the microbubbles PSF before localization^23^. In fact, applying adaptive beamforming without tissue canceling can lead to a distortion of the signal of microbubbles^45^. Here, the SVD was performed on a spatiotemporal matrix containing the raw IQ signals recorded by all channels before any processing. The Python code of this implementation is available in the provided git repertory^41^. The SVD threshold was empirically set. The raw RF data recorded by each piezoelectric channel were IQ-demodulated and then time-delayed with a voxel size equal to half the wavelength. A f-number of 1.5 was applied in both lateral and elevation directions. The different beamformers were computed once the tissue was filtered and the signal rephased. The value of *p* for pDAS varied from 2 to 4 and the number of iterations for iMAP from 1 to 4. The standard deviation for the Gaussian kernel in CFGF was fixed to one pixel as in^45^. The null voxels produced by the f-number were discarded to compute CF, CFGF, and iMAP.

Volumetric ULM post-processing was done with the same steps as described in^18^, briefly, the sub-wavelength localization step was done using radial symmetry^50^, and the tracking step was done using ’SimpleTracker’ with the Hungarian algorithm^51^. All the volumetric ULM process was done on MATLAB (The Mathworks, USA, 2021b).

## Results

### Adaptive beamformers sharpen the PSFs of the microbubbles in vitro

Various metrics are used to quantify the effect of the four different beamformers employed in this study. Fig. 1 presents a snapshot of the volumetric rendering of the microbubble PSFs in water for all proposed beamformers. The values for each beamformer are normalized to their respective maximum. It is evident that the 2DAS, 1 to 4 MAP beamformers (Fig. 1**b,c,d** and **e**) produce a more compact PSF compared to the DAS beamformer (Fig. 1**a**). No obvious effect of the CF and CFGF (Fig.1**g** and **h**) beamformers on the microbubble size compared to DAS. Fig. 2 summarizes the results found *in vitro*. First, the Full Width Half Maximum (FWHM) in the three directions, namely *x* (lateral), *y* (elevational), and *z* (depth), are shown in Fig. 2**a, b**, and **c**, respectively. Smaller PSF is remarked with a higher *’p’* value in the pDAS beamformer. A similar effect is seen for the iMAP, smaller PSF in the three directions for a higher value of *’i’*. A reduced FWHM value is also observed in the CF and CFGF beamformer compared to the standard DAS. However, CFGF has a higher value than CF. The Peak Side-to-Main Lobe Ratio (PSMR) is computed in Fig. 2**d**. A higher absolute value of the PSMR indicates fewer side lobes, hence higher contrast. Again the same effect is shown, with the increase of the *’p’* value in the pDAS and the *’i’* value in the iMAP beamformers, the higher absolute value of the PSMR in dB. It is also noticed that all the beamformers have fewer side lobes than the standard DAS beamformer. Finally, the signal-to-noise ratio (SNR) was measured on every PSF of the microbubbles *in vitro*, and a higher value of SNR was reached with all the beamformers compared to the standard DAS beamformer. Two to three-fold higher values can be seen with the higher values of *’p’* in the pDAS and *’i’* in the iMAP beamformers (Fig. 2**e**). The CF and CFGF had a higher value than DAS but intermediate between the highest value of *’p’* and *’i’* in pDAS and iMAP.

**Figure 2.**
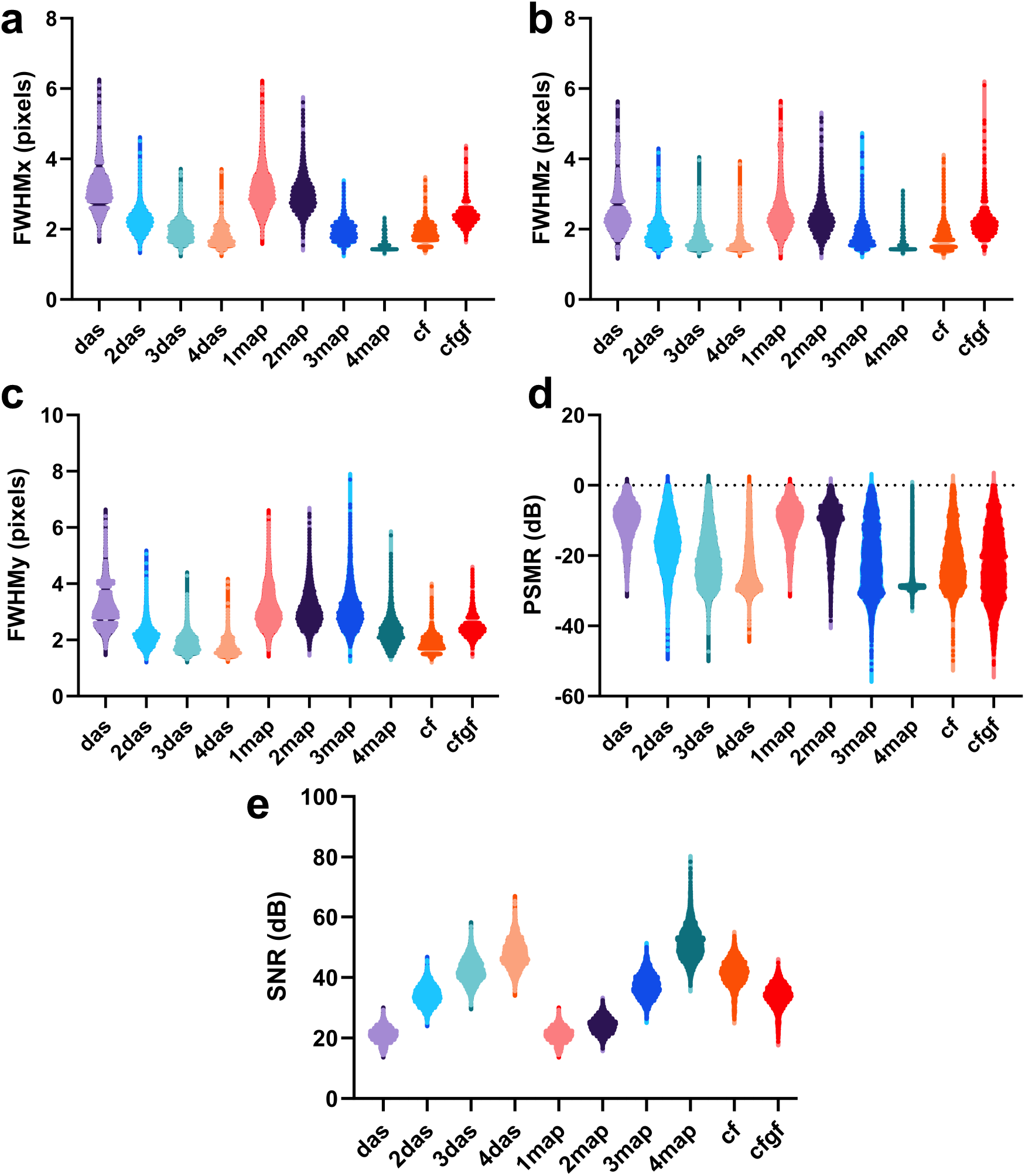
In-vitro adaptive beamforming metrics. Full Width Half Maximum (FWHM) in **a** x (lateral), **b** y (elevational) and **c** z (depth) direction. **d** Peak Side-to-Main Lobe Ratio (PSMR) and **e** Signal-to-Noise Ratio (SNR).

### Adaptive beamformer enhances volumetric ULM resolution in vivo

In Fig. 3, we represent the volumetric ULM rendering of the rat’s kidney with 8 different beamformers with the same color scale. For all the different adaptive beamformers, the following volumetric ULM parameters were used: the SVD cut-off was empirically determined and set to 8 out of 200. As in as in^18^, the detection was made with a local maximum of 900 MBs/volume with a local SNR set at 9 dB. The maximum linking distance was 0.23 mm, and the minimum length of accepted tracks was set to 15 frames (∼ 115 ms). In Fig. 3**a** represents volumetric ULM density for DAS beamformer, **b** 2DAS, **c** 1MAP, **d** 2MAP, **e** 3MAP, **f** 4MAP, **g** CF and **h** CFGF.

**Figure 3.**
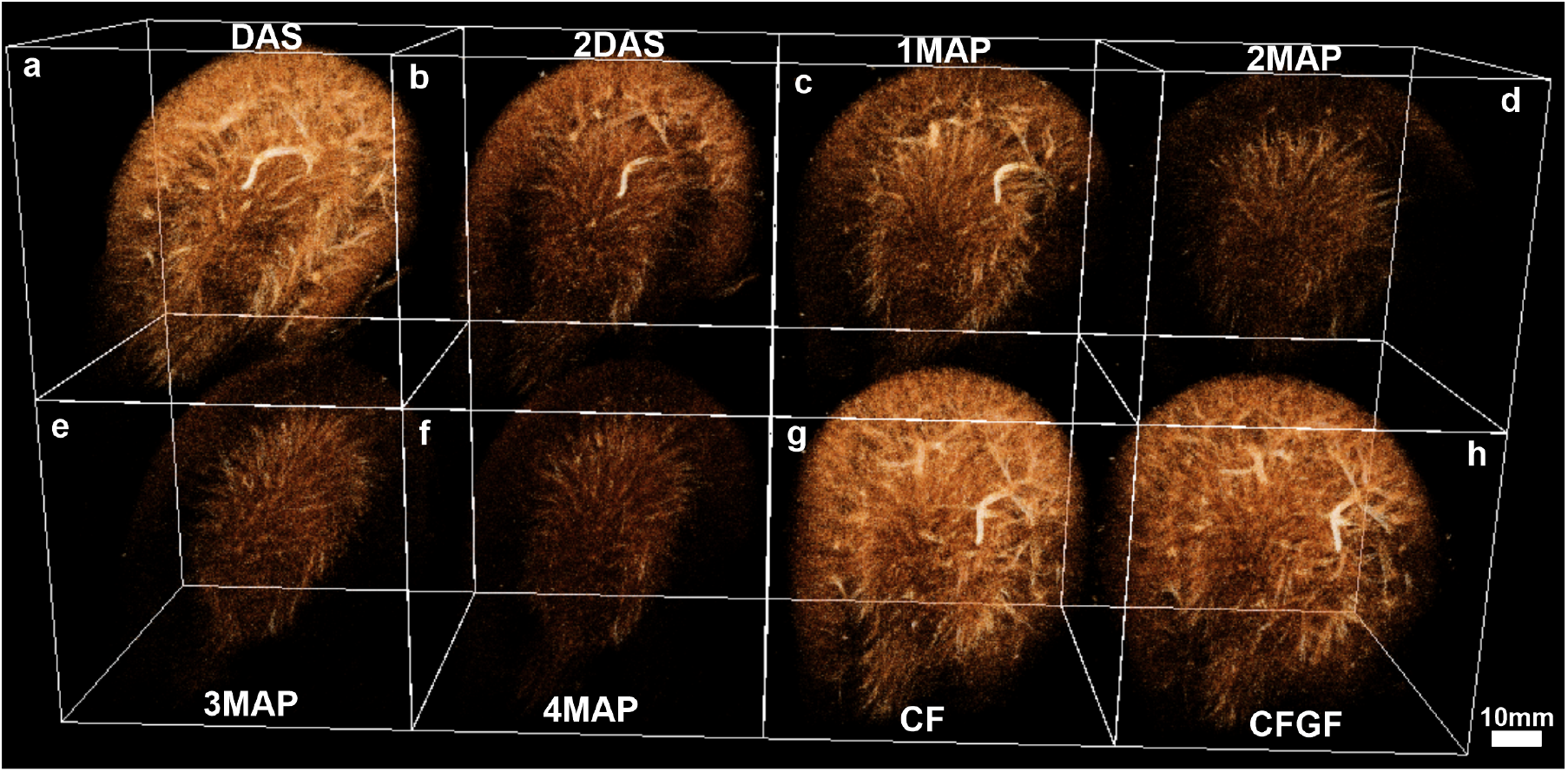
Volumetric ULM density map of the entire rat’s kidney for the proposed adaptive beamformers: **a** DAS, **b** 2DAS, **c** 1MAP, **d** 2MAP, **e** 3MAP, **f** 4MAP, **g** CF and **h** CFGF.

To quantify the effect of the adaptive beamformers on the final volumetric ULM images, several classical metrics were used in four different anatomical regions of the kidney, presented on Supplementary Fig. 7. As in^18^, a 3D region of interest (ROI) was designated within the kidney’s four principal regions: main vessels, inner medulla, outer medulla, and cortex. Each ROI was defined as a cubic structure with dimensions of 100 pixels per side, corresponding to 985 *μ*m. It is important to note that the cortical ROI was not visible from this perspective. In every part of the kidney, the track length has been measured for the different adaptive beamformers (see Fig. 4) on all the different regions of the kidney. In the main vessels (Fig. 4**a**), DAS, CF, and CFCG beamformers performed the best with longer tracks, followed by 2DAS and 1MAP right after, and finally, 2-4MAP performed the worst. In the upper left cortex of the kidney, (Fig. 4**b**) DAS, CF, and CFGF perfomed the best with a comparable performance of the 2DAS and 1MAP beamformers. Again, for this region, as in the main vessels, 2, 3, and 4MAP were inefficient in visualizing the cortex. For the inner and outer medulla (Fig. 4**c** and **d**, respectively), adaptive beamformers performed moderately better than the classical DAS. It is noted that higher-order beamformers 2DAS and 2MAP got the highest performance. Another way to look at the distribution of the track length metric is by computing the total tracks’ number for the different adaptive beamformers in the four different regions as seen in Fig. 5. We can see that in all the regions, DAS, CF, and CFGF beamformers scored the highest number of total tracks number. But in the inner medulla, the performance of particularly the 2DAS adaptive beamformer is better and comparable performance for the 1MAP in the outer medulla compared to the classical DAS beamformer.

**Figure 4.**
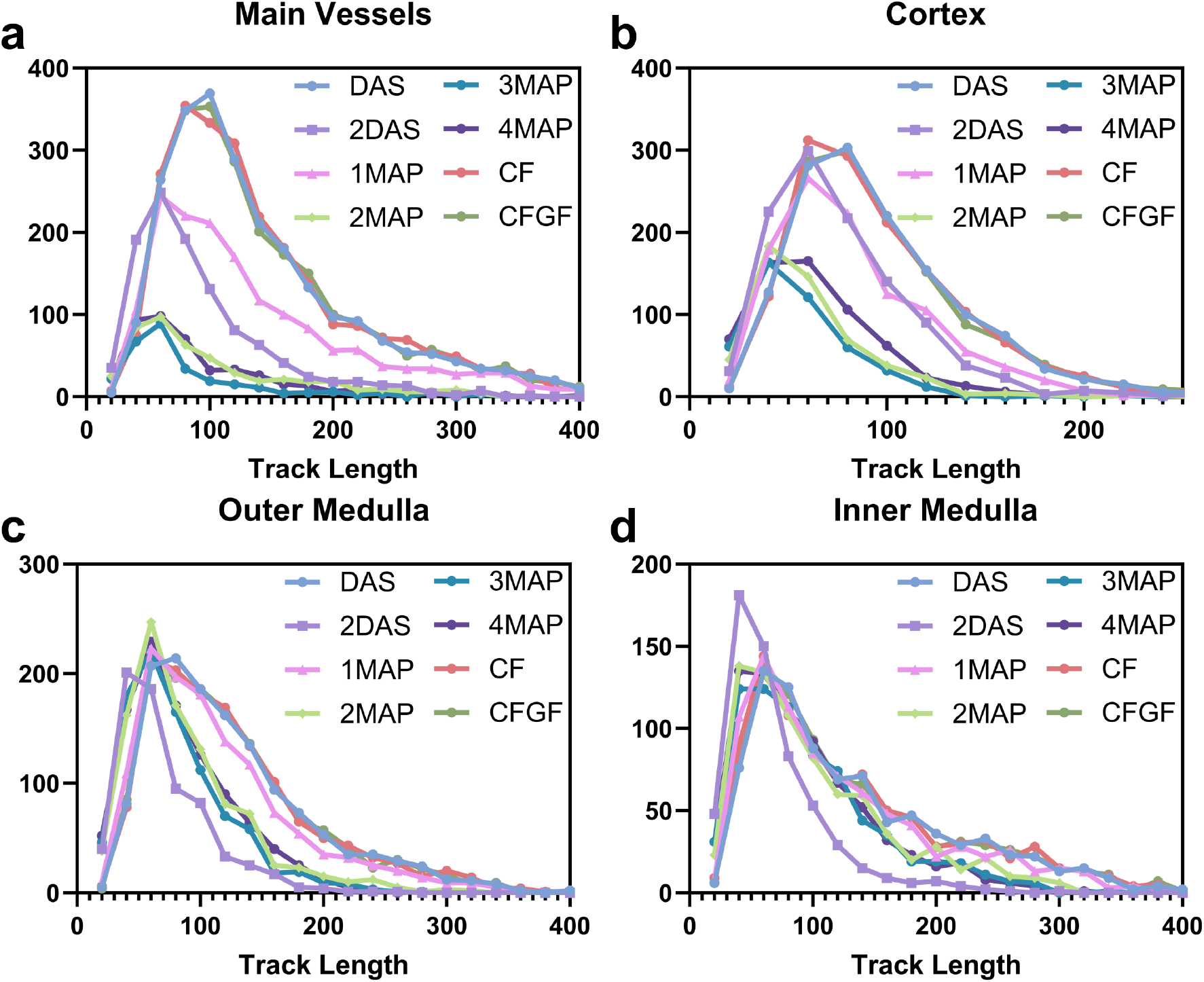
Volumetric ULM metric. Track length for the main kidney regions: **a** main vessels, **b** cortex, **c** outer medulla and **d** inner medulla.

**Figure 5.**
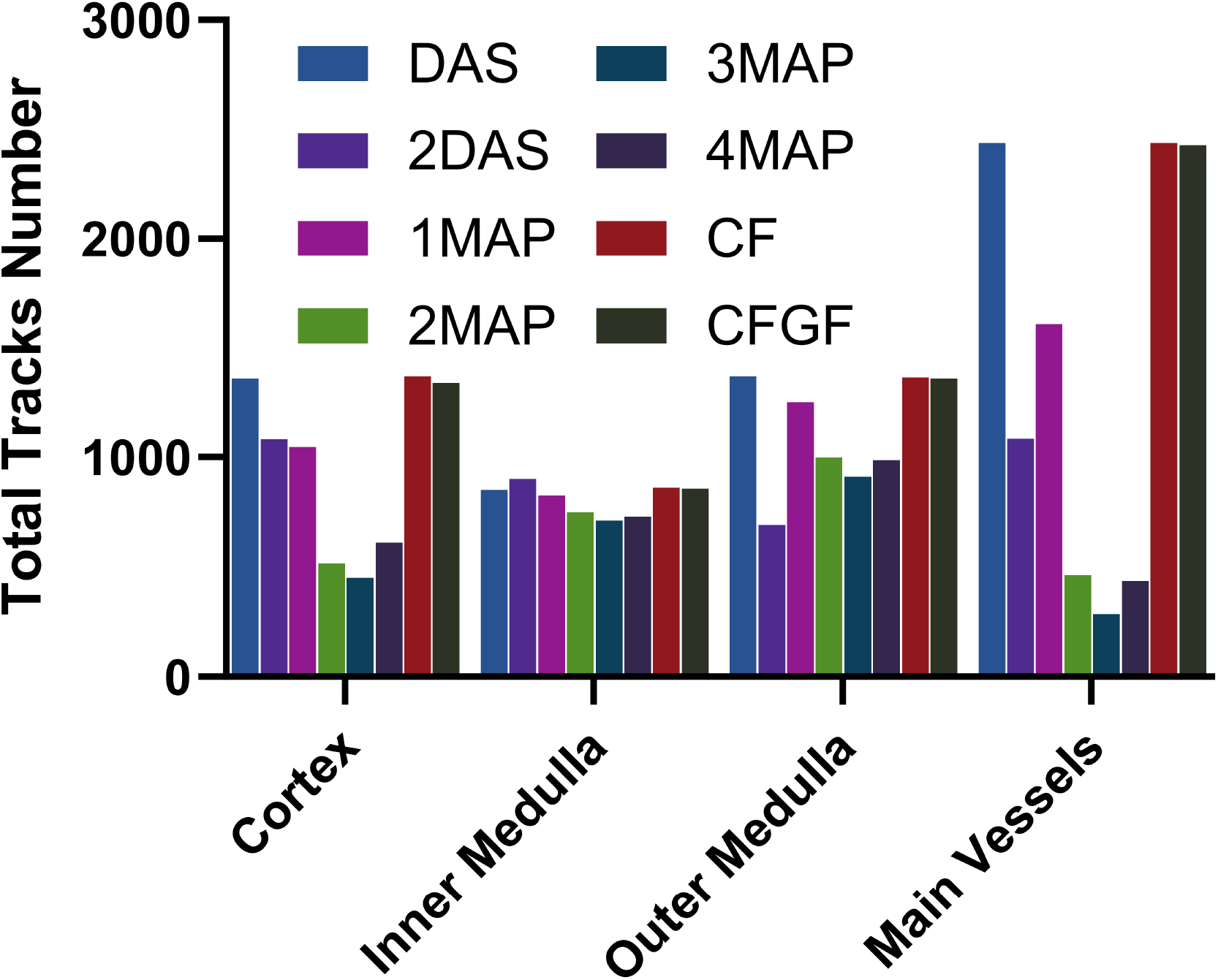
Volumetric ULM metric. Track counts for the main kidney regions: main vessels, cortex, outer medulla and inner medulla.

**Figure 6.**
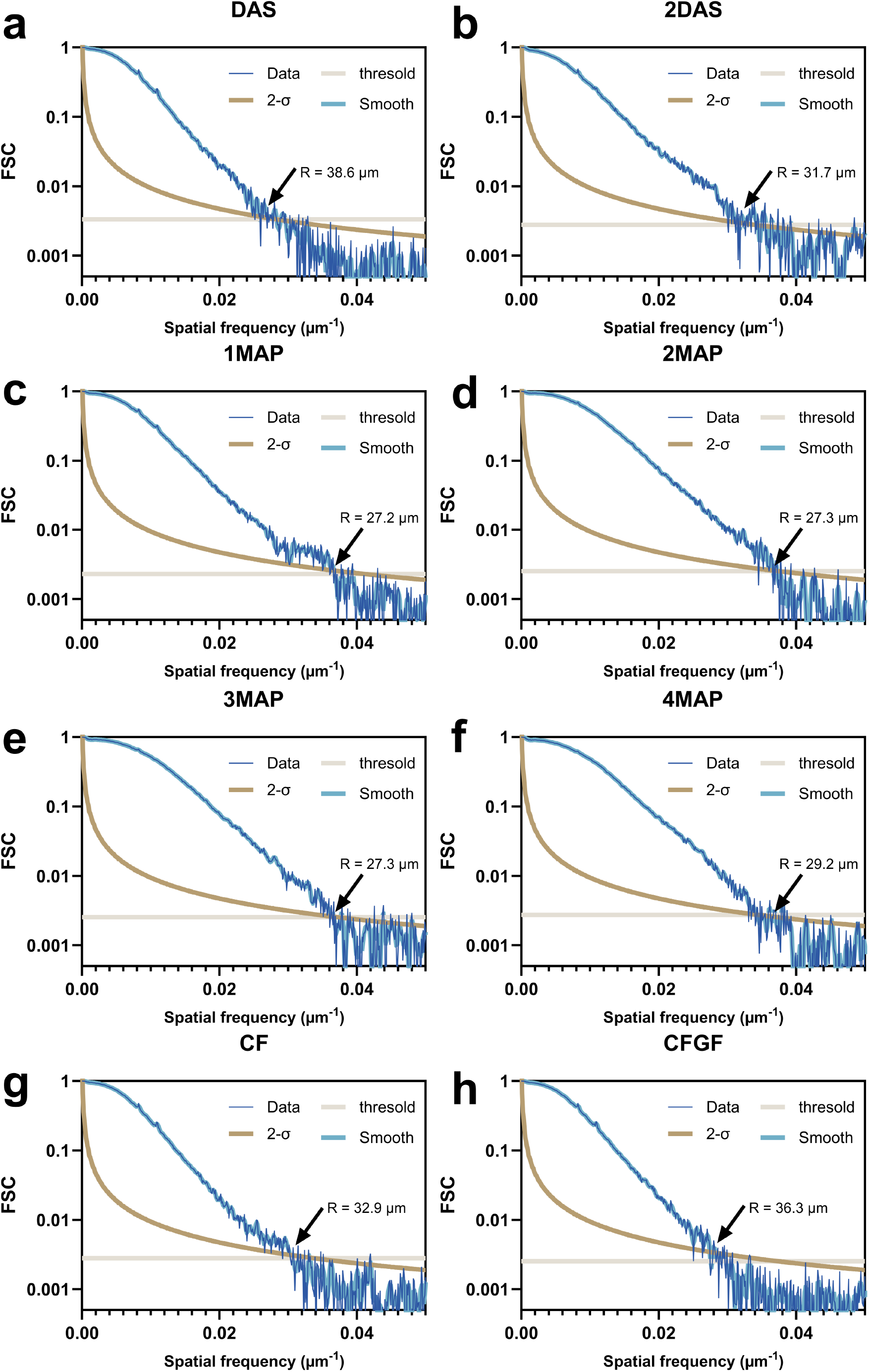
Volumetric ULM metric. Fourier Shell Correlation (FSC)^52^ for the proposed adaptive beamformers: **a** DAS, **b** 2DAS, **c** 1MAP, **d** 2MAP, **e** 3MAP, **f** 4MAP, **g** CF and **h** CFGF.

**Figure 7.**
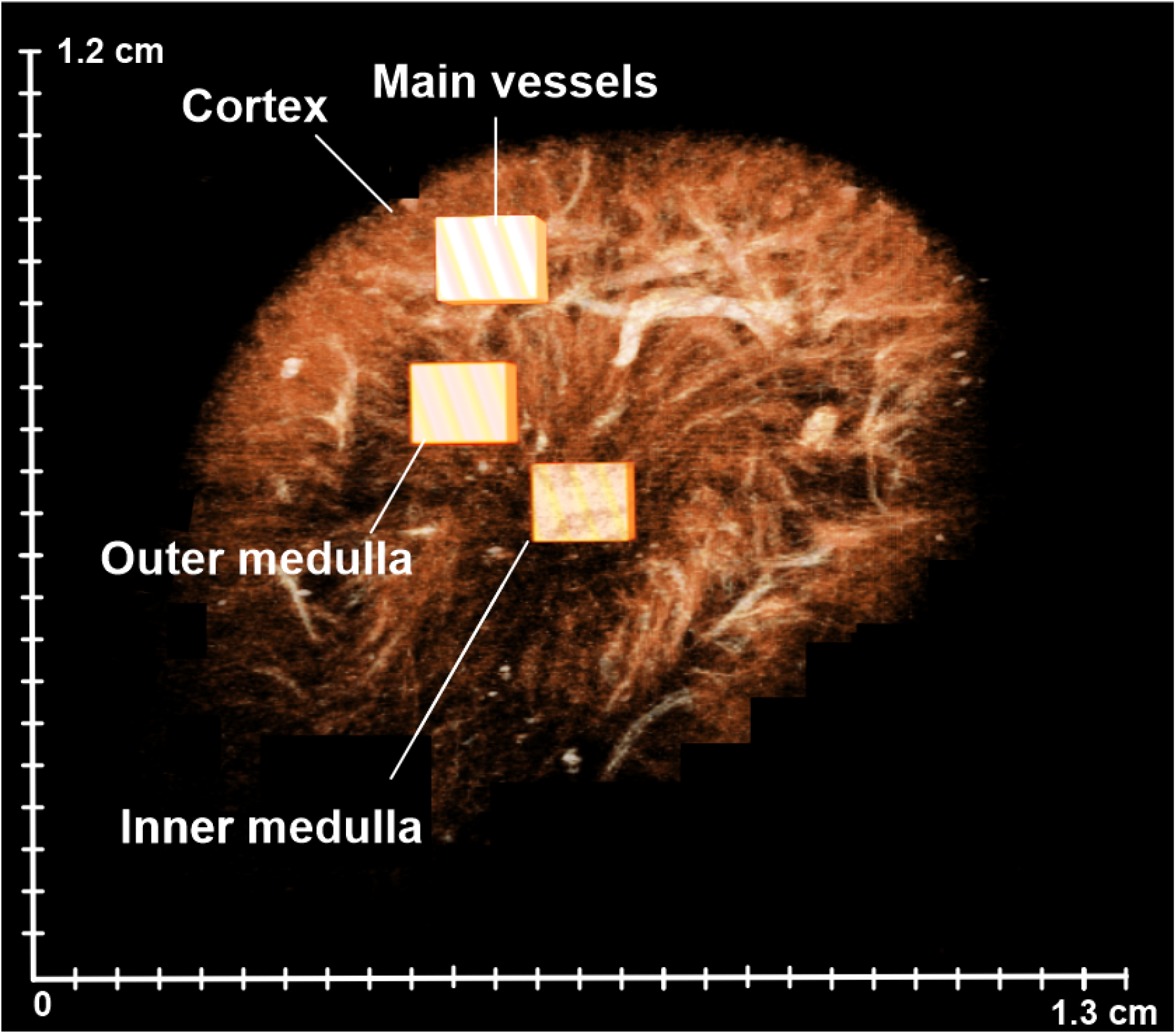
3D ROI of different renal zones were selected to quantify the adaptive beamforming effect on the volumetric ULM.

**Figure 8.**
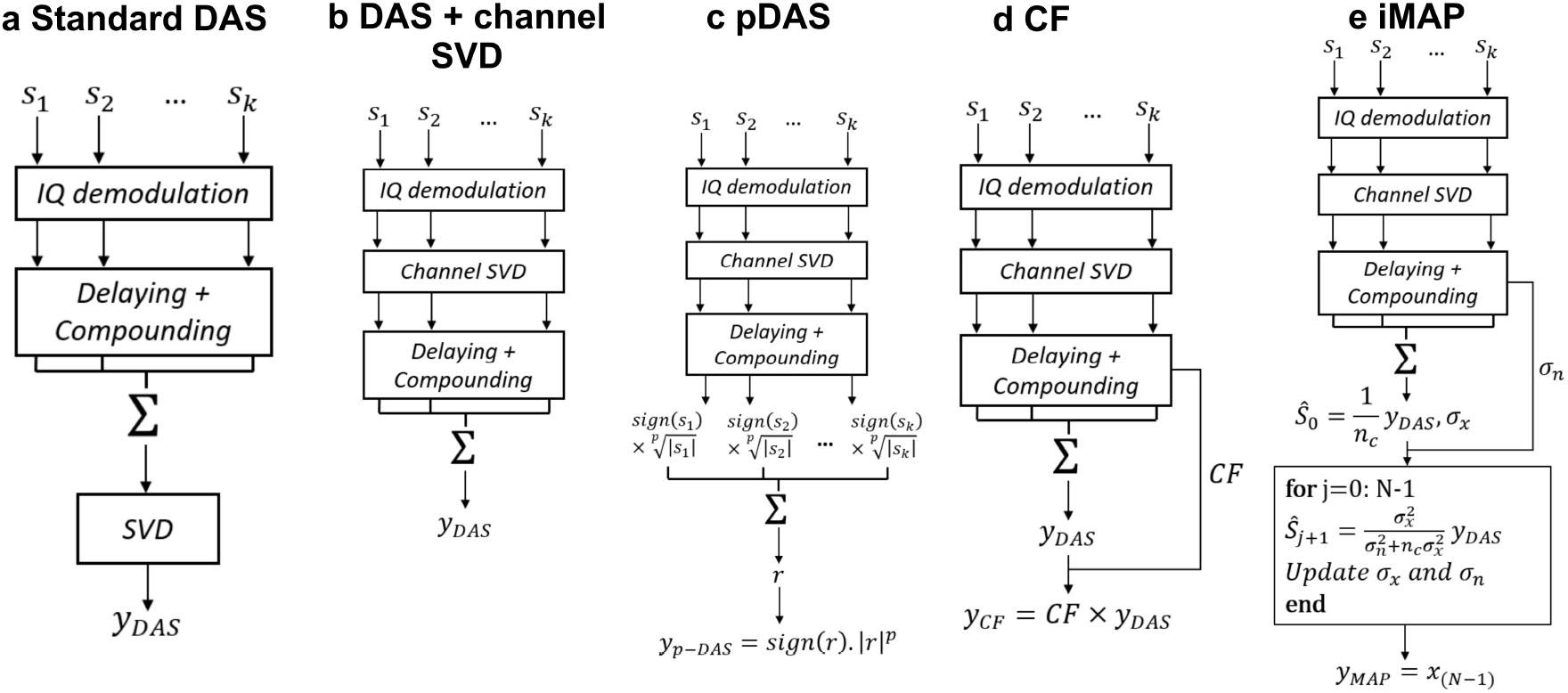
Graphical presentation of the beamformers and the clutter filtering: **a** standard DAS beamforming and tissue clutter filtering after beamforming, **b** standard DAS beamforming with channel-SVD before delaying and compounding. Different adaptive beamforming with channel-SVD such as **c** pDAS, **d** Coherence Factor, and **e** iMAP.

Finally, the last metric used in this work to quantify the effect of the adaptive beamformers is the classical metric used in ULM to measure the final resolution called the Fourier Shell Correlation (FSC)^52^. Fig. 6 shows the FSC of 8 different adaptive beamformers including **a** DAS with a resolution at 38.6 *μ*m, moderately higher resolution for the 2DAS beamformer in **b** at 31.7 *μ*m as well as for the CF and CFGF beamformers in **g** and **h** at 32.9 *μ*m and 36.3 *μ*m, respectively. The iMAP beamformers had a moderately better resolution ranging from 27.2 *μ*m **c**, 27.3 *μ*m for both 2MAP and 3MAP in **d** and **e**, respectively, and 29.2 *μ*m for the 4MAP beamformer in **f**. Note that the values of FSC should be taken into consideration depending on the total number of tracks as it will be discussed in the next section.

## Discussion

Different adaptive beamformers were employed to study their effect on volumetric ULM. It is important to emphasize that the tissue clutter filtering was done using a channel-SVD strategy before the beamforming step. With this process, the impact of the adaptive beamformers was directly on the microbubble PSF. When checking the effect *in vitro*, the size of the PSFs was generally reduced with an increased adaptive beamformer order (Fig. 2**a,b** and **c**). Also, a higher order of adaptivity improved the resolution by lowering the size of the side-lobes, as seen with the PSMR value getting higher (in absolute value, Fig. 2**c**. Finally, the SNR of the microbubbles is also significantly higher with the order of adaptive beamformers going from 20 dB to 40 dB with 4DAS and 4MAP. The *in vitro* results agreed with the previous findings reported by Yan *et al*.^23^. The microvessel architecture of the kidney is shown on the volumetric ULM density rendering in Fig. 3 with the eight proposed beamformers. As we can see, higher orders of the adaptive beamforming led to a restricted view of the kidney vasculature (Fig. 3**b-f**). This might be due to the high coherence in space and time of the microbubbles moving inside the medulla. As seen in our previous study^18^, the flow speed in the inner and outer medulla was the lowest among the different regions, and the trajectory length was the highest. Qualitatively, the CF and CFGF beamformers showed a comparable good image as the classical DAS beamformer.

To precisely illustrate the effect of the proposed beamformers, four different ROIs were selected within the kidney: main vessels, cortex, inner medulla, and outer medulla. The microbubble track length and the total number of tracks wer measured in each region. Generally, increased adaptivity resulted in shorter track lengths (Fig. 4) and lower number of tracks (Fig. 5). As stated, high-order adaptive beamformers enhance the medulla part where the microbubbles had a long and slow path. 2DAS and 2-4MAP had a comparable number of tracks as standard DAS and even higher in the inner medulla for the 2DAS (Fig. 4 and Fig. 5). Finally, CF and CFGF were moderately better than the DAS beamformer in all kidney regions.

The last metric used was the Fourier Shell Correlation (FSC), now often performed in ULM characterization. Careful attention needs to be taken before concluding with the FSC. FSC separates the tracks of the image into two subsets randomly and then applies the correlation in the Fourier domain. A threshold is then set to determine the smallest reproducible spatial scale between the two sets^31^. In some cases, a reduced number of tracks following common paths would create an artefactual enhanced resolution, which is inappropriate. For instance, in Fig. 7, 2-4MAP had the highest resolution around 20 *μ*m while the whole kidney part was not fully reconstructed on the image. On the other hand, 2DAS and 1MAP performed equally with relatively close resolution but with fewer tracks (Fig. 6**b** and **c**). Finally, we notice that the CF beamformer has an equivalent number of tracks to the standard DAS beamformer but with a significantly higher resolution.

## Conclusion

In conclusion, we studied the effect of multiple adaptive beamformers on the final volumetric super-resolution images of the rat’s kidney. Channel-SVD was applied before beamforming to directly influence the microbubble PSF. We demonstrated that the Coherence Factor (CF) beamformer can moderately enhance the final resolution. In contrast, higher-order adaptive beamformers such as 2DAS and 2-4 MAP can significantly improve the medulla region of the kidney. Additionally, we advocate for open science. We have made a GPU-accelerated 3D beamformer with and without adaptive beamforming code available online, along with the *in vivo* data used for this study.

## Acknowledgment

This work was partly supported by the European Research Council (ERC) through the European Union Horizon 586 H2020 Program/ERC Consolidator under Grant 772786-ResolveStroke. This work was supported by the LABEX PRIMES (ANR-11-LABX-0063) and LABEX CELYA (ANR-10-LABX-0060) of the Université de Lyon, within the program “Investissements d’Avenir” (ANR-11-IDEX-0007), operated by the French National Research Agency (ANR). It was also partly funded by France Life Imaging from the French “Investissements d’Avenir” program (ANR-11-INBS-0006).

